# Arabidopsis lateral shoots display two distinct phases of growth angle control

**DOI:** 10.1101/2023.03.31.535051

**Authors:** Martina De Angelis, Stefan Kepinski

**Affiliations:** Centre for Plant Sciences, University of Leeds, UK

## Abstract

Shoot growth angle is a fundamental determinant of plant form. In their later development, lateral branches maintain gravitropic setpoint angles (GSAs) in which growth is set and maintained relative to gravity. The typically non-vertical GSAs are the product of an auxin-dependent antigravitropic offset that counteracts underlying gravitropic response in the branch (Roychoudhry *et al*., 2013). Here we describe an earlier phase of branch development in which the young lateral shoot grows rootward, independently of gravity, promoting a spreading growth habit. In normal development, this phase of growth is terminated with the onset of the GSA programme, with branches then growing upwards to assume their mature form. The biophysical basis of the early rootward phase of branch growth can be traced back to greater cell proliferation on the upper, adaxial side that upon expansion, drives asymmetric growth. Our data indicate that cytokinin is involved in this process and that the transcription factor TCP1 is an important regulator of lateral shoot adaxial identity and differential ad-abaxial cell proliferation.

## Introduction

Since the colonisation of land, plants have evolved a terrific number of shapes both above and below ground, with shoot branching contributing significantly to this diversity. In Arabidopsis, lateral shoots originate from axillary meristems (AMs) also referred as axillary buds, which are initiated from a population of undifferentiated cells in the leaf axil expressing the meristematic marker *SHOOT MERISTEMLESS* (*STM*) (Wang and Jiao, 2018). Up-regulation of *STM* determines AM initiation through the activation of cell division (Shi *et al*., 2016). This is followed by *de novo* expression of *WUSCHEL* (*WUS*) and the establishment of a new organization centre with the same potential as the shoot apical meristem (Wang *et al*., 2017; Xin *et al*., 2017). Low levels of auxin are required in the leaf axil to maintain the meristematic competence of AM precursors and allow *STM* up-regulation, while cytokinin (CK) signalling activates *WUS* expression and contributes to maintain the stem-cell niche organised (Wang *et al*., 2014). In addition, CK positively regulates the cell cycle through the transcriptional activation of type-D3 cyclins (CYCD3), which in turn are responsible for triggering the G1/S transition in dividing cells (Riou-Khamlichi *et al*., 1999; Dewitte *et al*., 2007). After its initiation, the AM can either remain dormant or outgrow into a lateral shoot, depending on both endogenous and exogenous factors, such as hormones, light and nutrients (Wang *et al*., 2019).

After bud outgrowth, lateral shoots are maintained at specific angles with respect to gravity, independently of other parts of the plant. These growth angles have first been characterised by Digby and Firn and named gravitropic setpoint angles (GSAs) (Digby and Firn, 1995). According to the GSA concept, organs that are maintained at the vertical, such as primary roots and shoots, have GSAs of 0° and 180°, respectively, while lateral shoots and roots maintain non-vertical GSAs between these two extremes. The perception of gravity in Arabidopsis primary and lateral shoots occurs in the endodermis layer, in which dense, starch-filled plastids sediment in response to gravity and function as statoliths (Fukaki and Tasaka, 1999; Chauvet *et al*., 2016). In primary shoots, gravitropism is responsible for the maintenance of the vertical GSA (Went, 1926; Cholodny, 1927). Reorientation of the primary shoot away from the vertical drives statolith re-sedimentation within endodermal cells, leading to the PIN protein-mediated redistribution of auxin to the lower side of the stem (Ottenschläger *et al*., 2003). Here, auxin promotes cell expansion inducing asymmetric growth between the upper and lower side of the stem, that allows the primary shoot to bend upward and restore its vertical GSA (Went, 1926; Cholodny, 1927; Rayle and Cleland, 1980). In non-vertically growing lateral shoots, the action of gravitropism, which would otherwise move the branch to the vertical, has been shown to be counteracted by an auxin transport-dependent antigravitropic offset (AGO) mechanism that permits stable non-vertical growth (Roychoudhry *et al*., 2013). The AGO mechanism has been further characterised in lateral roots, where it has been shown to act at the level of PIN-FORMED (PIN) auxin efflux carrier polarisation in gravity-sensing cells such that gravitropic auxin transport to the lower side of the lateral root growing at GSA is balanced by auxin exported to the upper side via the AGO (Roychoudhry *et al*., 2013, 2019). Although the molecular basis of non-vertical GSA maintenance has been mainly studied in lateral roots, analysis of GSA-impaired mutants suggests that similar molecular mechanisms are involved in the control of lateral shoot GSAs (Roychoudhry *et al*., 2013; Yoshihara, Spalding and Iino, 2013). In addition to PIN protein regulation, this control involves the action of members of LAZY-LIKE (LZY) family, which have been proposed to influence PIN polarity in gravity sensing cells (Taniguchi *et al*., 2017; Kawamoto *et al*., 2020). In lateral shoots *LZY1, LZY2* and *LZY3* are redundantly expressed in the endodermis, with *LZY1* loss of function showing the stronger defects in GSA control and gravitropism among the single mutants (Yoshihara, Spalding and Iino, 2013; Taniguchi *et al*., 2017).

Studies on lateral shoot development have focused on axillary bud formation/activation and mature branch regulation, leaving the early development of growth angle unexamined. Here we describe a novel phase of lateral shoot development, before the onset of GSA maintenance, in which branches grow rootward. We demonstrate that young lateral shoots, although gravicompetent, do not maintain growth angles as GSAs and indeed, their development is independent of gravity. We show that this pattern of early branch growth can be traced back to a difference in cell number between the adaxial and the abaxial side of young lateral shoots, which upon cell expansion, translates into the observed rootward bending of young lateral shoots. Our data also indicate that cytokinins and the transcription factor TCP1 are involved in the specification of ad-abaxial polarity in lateral shoots.

## Materials and Methods

### Plant material and growth conditions

*Arabidopsis thaliana* lines used were in the Col-0 background. The *pgm1* (Caspar and Pickard, 1989), *lzy1* (Yoshihara, Spalding and Iino, 2013), *scr-3* (Fukaki *et al*., 1998), *CYCD3;1-3::GUS* (Dewitte *et al*., 2007), *ahk2-2ahk3-3* [*ahk2ahk3*], *ahk2-2cre1-12* [*ahk2ahk4*], *ahk3-3cre1-12* [*ahk3ahk4*] (Higuchi *et al*., 2004), *rock2* and *rock3* (Bartrina *et al*., 2017), *ipt3-2ipt5-2ipt7-1 [ipt3ipt5ipt7]* (Miyawaki *et al*., 2006) lines have been previously described. *p35S::TCP1* line was obtained by cloning *TCP1* CDS into pALLIGATORIII (derived from *pFP101* (Pan *et al*., 2001)) using Gateway system. Seeds were sown in soil in 52×50 mm pots and stratified at 4°C for two days. Plants were grown in glasshouses or growth rooms at 20-22°C under long day photoperiod (16 h day).

### Clinorotation experiment

Plants at rosette stage (~ 2-3 weeks old) were placed on a 1-D clinostat with the axis of rotation parallel to the shoot-root axis of the plant, while control plants were left clinorotating vertically. A Mikrops Electric Clinostat (Flatters & Garnett, Manchester, UK) at 4 rph (rotations per hour) was used. Plants were left clinorotating up to 2-3 weeks and photographs were taken after bolting following lateral shoot development for a week.

### Lateral shoot kinetics and reorientation assays

For lateral shoot kinetics, plants were grown in glasshouses or growth rooms for ~4 weeks. Once bolted plants were photographed every 24 hours following lateral shoot development. For the 45° reorientation assay, plants with stage III or stage IV lateral shoots were tilted 45° from the vertical, reorienting lateral shoots to lower angles with respect to the gravity vector. For the gravicompetence assay, plants were laid down on a side, reorienting lateral shoots perpendicularly to the gravity vector. To stop the primary shoot gravitropic response from affecting lateral branch inclination, in both assays primary shoots were secured to thin wooden sticks inserted in the pots following the primary axis of the plant. For reorientation assays performed in the dark, plants were kept in darkness for 1 hour prior to reorientation. Plants were imaged after 6 hours post-reorientation.

### Cell length and number measurements

Sections of plants containing a SII/SIII lateral shoot with the apical stem and meristem intact, were kept in tubes maintaining the natural branch orientation while getting fixed under vacuum with 4% Paraformaldehyde (PFA) in Phosphate-buffered saline (PBS) solution at pH 6.9 for 2 hours. Afterwards samples were washed twice in PBS and lateral shoots sectioned dividing the adaxial from the abaxial side of the branch. Sections were incubated with calcofluor white overnight and washed for at least 30 minutes in PBS before being imaged. Images were taken using a Zeiss LSM 880. Cellular length and number were measured using the software Fiji.

### GUS staining

Section of shoot growing lateral shoots at different developmental stages were fixed in ice with acetone 90% for 40 minutes. After that, the sections were washed twice in 100 mM phosphate buffer at pH 7 (Na_2_HPO_4_ + NaH_2_PO_4_) and incubated in Gus solution (100 mM phosphate buffer, 0.5 mM K_3_Fe(CN)_6_, 0.5 mM K_4_Fe(CN)_6_, 0.5-1 mg/ml X-Gluc dissolved in DMSO, 0.1% Triton X-100). Samples were then vacuumed for 10 minutes and left at 37°C overnight. Samples were then washed in 100% ethanol and cleared using a 3:1 solution of methanol and acetic acid. Samples were stored in 70% ethanol. Images were taken using a Zeiss Scope A1 microscope with an Axiocam 305 colour camera.

### Real-Time quantitative PCR

Quantification of *TCP1* mRNA expression levels were carried out in relation to *ELONGATION FACTOR 1α* (*EF1α*) and *ACTIN2* (*ACT2*) mRNAs using the geometric mean as described by (Taylor *et al*., 2019). The following primers were used in the reaction: *TCP1* 5’-TCTTCACTCTCTGGCCATCA −3’ and 5’-CTGCTGATACCATATGGCCC −3’, *EF1α* 5’-CTTCAAGTACGCATGGGTGT −3’ and 5’-CTTGGTGGTCTCGAACTTCC −3’, *ACT2* 5’-AATTTCCCGCTCTGCTGTT −3’ and 5’-TGCCAATCTACGAGGGTTTCT −3’.

## Results

### Young lateral shoots are gravicompetent

After bud outgrowth, we observed that newly formed Arabidopsis lateral shoots elongate, growing away from the primary shoot and assuming a rootward trajectory (Fig. 1A). This first period of growth lasts 5 to 7 days and precedes the setting of the GSA, indicating the existence of a new phase of lateral shoot development that has not yet been described. To characterise this early phase in more detail, we divided lateral shoot development into four stages, based on the changes in growth angle shown by the branch (Fig. 1A). During stage I (SI) lateral shoots elongate vertically, maintaining proximity with the primary stem. This phase is followed by a rootward bending stage (Stage II/ SII) that ends after 2 to 3 days, with lateral shoots reaching a nearly horizontal growth angle (Stage III/ SIII). Stage II to III rootward growth is followed by lateral shoots bending shootward and setting their given GSA, that will be actively maintained for the rest of the plant life (Stage IV/ SIV) (Roychoudhry *et al*., 2013). To understand the nature of SII-SIII rootward bending and the role played by gravity in the process, we performed two different reorientation assays to discriminate GSA maintenance from a more general gravicompetence in early developmental stages. First, we tested the capacity of young lateral shoots to maintain GSAs tilting plants 45° from the vertical, so that branches were reoriented to lower angles (Fig. S1). As previously demonstrated, after 6 hours SIV lateral shoots bend upwards restoring their previous GSA (Roychoudhry *et al*., 2013). SII-III lateral shoots, instead, do not respond to this gravistimulation and do not alter their growth in response to the new position they are placed (Fig. 1B).

**Figure 1:**
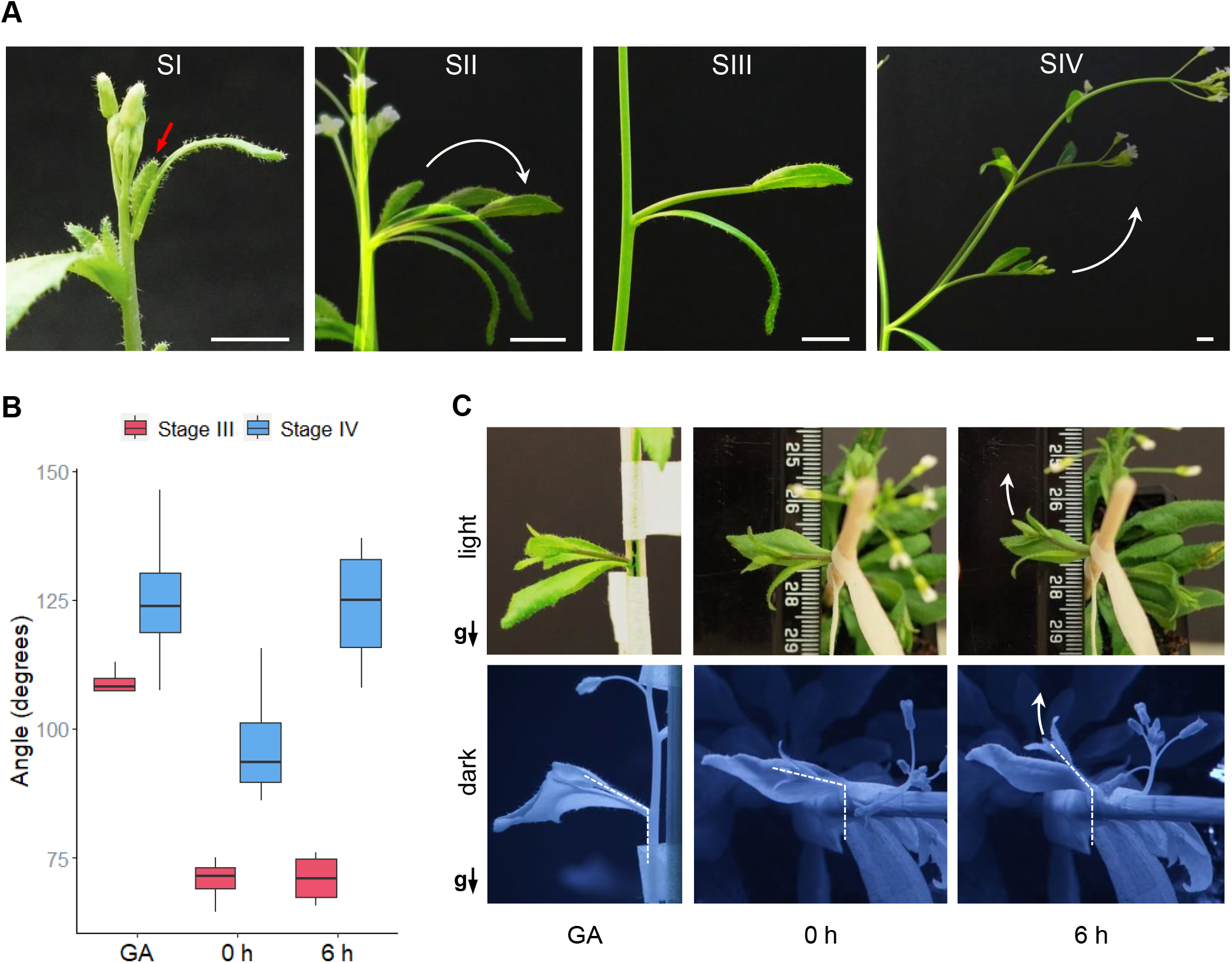
Early developmental stages of lateral shoots in Arabidopsis are gravicompetent. A) Representative images of the four developmental stages of Arabidopsis lateral shoots. SII and SIV images are obtained overlaying three pictures taken 24 hours from each other. Red arrow indicates the SI lateral shoot, white arrows indicate the direction of the lateral shoot growth. Scale bars = 0.5 cm. B) Quantitative analysis of 45° reorientation of SIII and SIV lateral shoots. GA = Growth Angle, 0 h = 0 hours after reorientation, 6 h = 6 hours after reorientation. C) Representative images of SII-SIII lateral shoots during “perpendicular assay” performed in the light or in the dark. Black arrows indicate the direction of gravity, g = gravity. White arrows indicate the direction of the bending, dashed lines indicate the angle of lateral shoot during the reorientation with respect to gravity.

A second type of reorientation assay was devised to test if, although not responding to the 45° tilting assay, young lateral shoots still show gravicompetence (Fig. S1). Plants were laid horizontally, and lateral shoots reoriented perpendicular to the gravity vector. For this reason, we called this assay “perpendicular assay”. 6 hours following the perpendicular reorientation, both SII and SIII lateral shoots bent upwards (Fig. 1C). This behaviour was observed both in the light and in the dark (Fig. 1C), confirming that, although not maintaining GSAs, lateral shoots have the capacity to sense and respond to gravity. Further, any contribution from phototropism to the bending response can be excluded.

### Early stages of lateral shoot development are independent of gravity

Since young lateral shoots can perceive and respond to gravity, we used clinorotation to understand the contribution of gravity to the early stages of lateral shoot development. Clinorotation induces an omnilateral gravitational stimulus to the plant that prevents the statoliths from sedimenting in the cell and determining the gravity direction. This method has been used to good effect to remove a stable reference to gravity during plant growth (Lyon and Yokoyama, 1966; Heathcote, Chapman and Brown, 1995; Roychoudhry *et al*., 2013). Plants were mounted on a clinostat before bolting and subjected to clinorotation for the entire duration of development of the primary and lateral shoots, meaning that lateral shoots had never experienced a stable, polarising gravity reference. Under clinorotation, lateral shoot development progresses from stage I to stage II similarly to control plants (Fig. 2A). After that, lateral shoots continue to bend rootward without entering SIII and SIV, while non-clinorotated control plants stop bending rootward in SIII and set GSA normally in SIV (Fig. 2B-C). This suggests the existence of two distinct phases in lateral shoot development: an early gravity-independent phase consisting of SI and SII, and a following gravity-dependent phase that start with the arrest of lateral shoot rootward growth in SIII, and terminates setting the GSA in SIV.

**Figure 2:**
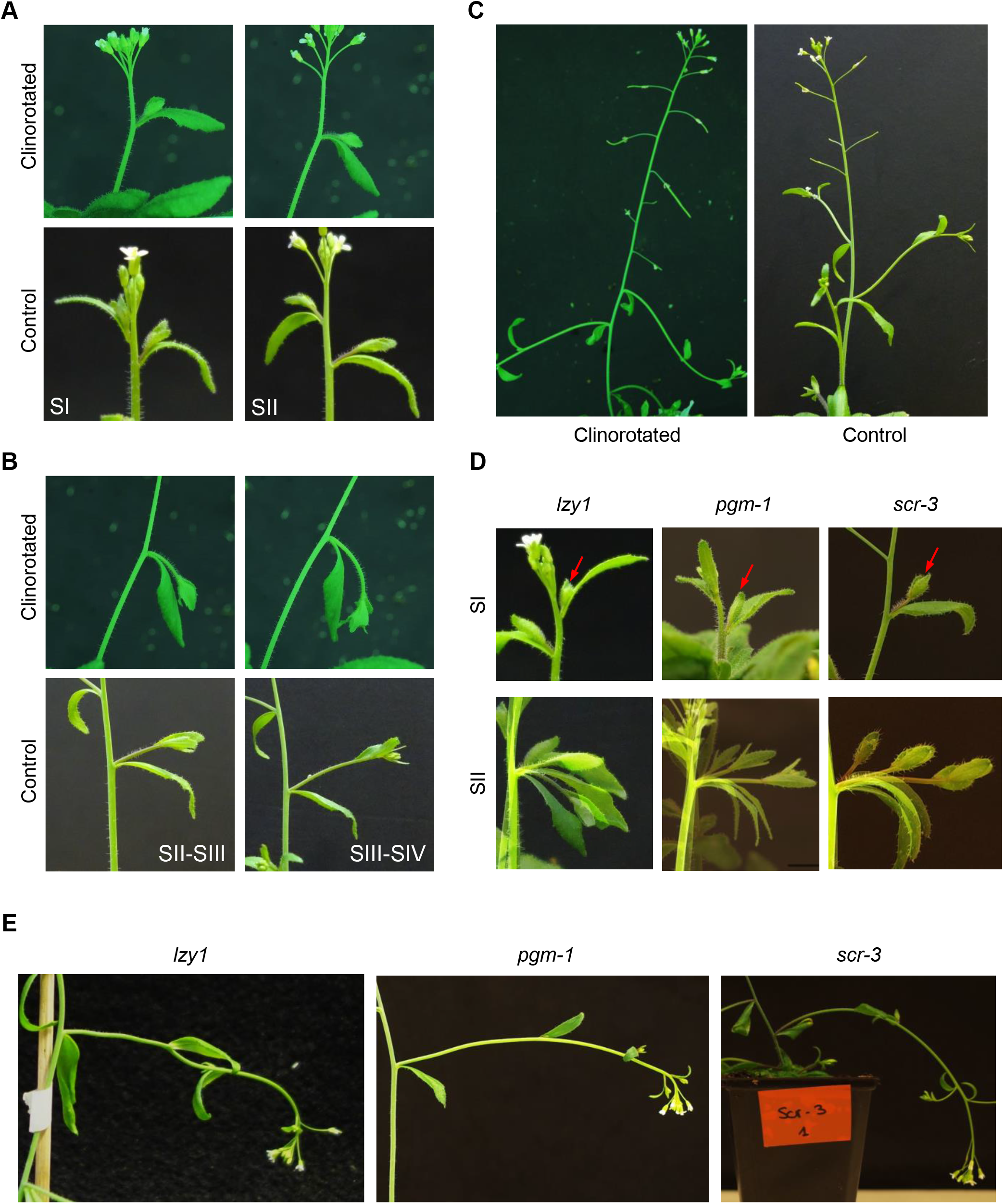
SI and SII are set independently of gravity. A-B) Representative images of lateral shoot early developmental stages (SI-SII) of clinorotated and non-clinorotated plants (control). C-D) Representative images of late developmental stages (SIII-SIV) of lateral shoots during clinorotation and in control plants. D) Representative images of SI and SII lateral shoots of *Izy, pgm-1* and *scr-3* mutants. SII images are obtained overlaying pictures taken every 24 hours from each other. Red arrows indicate the lateral shoot. E) Representative images of SIV lateral shoots of *lzy1, pgm-1* and *scr-3* mutants.

To confirm this hypothesis, we followed lateral shoot development of *PHOSPHOGLUCOMUTASE* (*PGM*), *SCARECROW* (*SCR*) and *LZY1* loss-of-function mutants *pgm-1, scr-3* and *lzy1* (Caspar and Pickard, 1989; Fukaki *et al*., 1998; Yoshihara, Spalding and Iino, 2013). Both *pgm-1* and *scr-3* have gravity-sensing impairments due to lack of starch-filled statoliths and endodermis layer respectively (Caspar and Pickard, 1989; Fukaki *et al*., 1998). As previously demonstrated, the three mutants failed to maintain normal GSAs in SIV lateral shoots (Fig. 2E). Conversely early SI, and SII lateral shoots in these mutants do not show any significant alteration and are comparable with those of WT (Fig. 2D). This shows that although *pgm-1, scr-3* and *lzy1* do not maintain SIV GSAs, lateral shoot early development is not impaired and can be considered independent from the maintenance of GSA.

### Lateral shoot adaxial and abaxial sides have distinctly different numbers of cells

In order to understand the biophysical basis of rootward bending in young lateral branches, we looked for differences in cell length and/or cell number between the adaxial and the abaxial side of the shoot that might explain the rapid change in growth angle displayed during SII rootward growth leading to the nearly horizontal growth angle in SIII. As expected across the lateral shoot, epidermis cells show a proximal-distal polarity with respect to the primary shoot, with distal cells, closer to the lateral floral meristem, having meristematic morphology and increasing their size and differentiation towards the proximal region of the branch (Fig. 3A). We decided to measure cell length and cell number of the distal region of SIII lateral shoots (Fig. 3A). Here, no significant differences were found in epidermal cell length between the adaxial and abaxial side (Fig. 3B), while in the same area, the adaxial side contained around 21% (± 5%) more cells per 0.1 mm^2^ than the abaxial side (Fig. 3B). To confirm the differences in cell number, we also analysed cell number in the first layer of cortical cells immediately under the epidermis. As in the epidermis, cortical cells on the adaxial side contained approximately 17% (± 4%) more cells per mm compared to the abaxial side (Fig. 3B).

**Figure 3:**
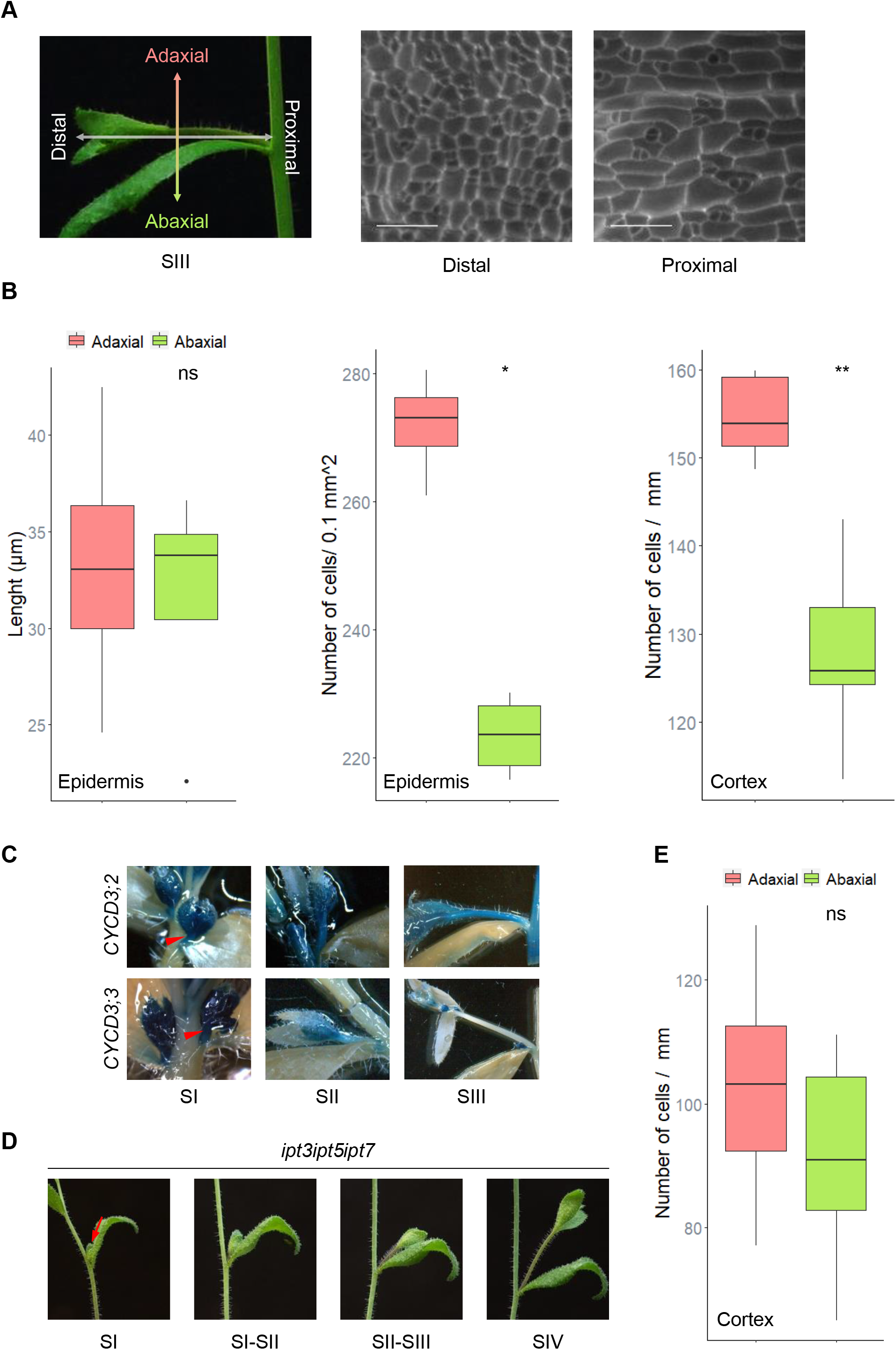
Adaxial and abaxial side of young lateral shoots show differences in cell proliferation. A) Representative images of a SIII lateral shoot with marked ad-abaxial and proximal-distal polarities, and epidermis layer of lateral shoot proximal and distal region. Scale bar = 50 μm. B) Quantitative analysis of cell length (epidermis) and number (epidermis and cortex) in the distal region of WT SII-III lateral shoots. C) Representative images of *CYCD3;2::GUS* and *CYCD3;3::GUS* reporter lines during lateral shoot development. Red arrows indicate lateral shoots at SI. D) Representative images of lateral shoot development of *ipt3ipt5ipt7*. Red arrow indicates SI lateral shoot. E) Quantitative analysis of *ipt3ipt5ipt7* cell number in lateral shoot cortex. Statistical analyses were performed using Wilcoxon test. ns = non significant, (*) = p < 0.05, (**) = p < 0.005.

In the shoot, cytokinins (CK) are involved in activating cell division through the regulation of D3 cyclins (CYCD3) (Riou-Khamlichi *et al*., 1999). In Arabidopsis, there are three *CYCD3* genes with partially overlapping expression, controlling cell proliferation during leaf growth (Dewitte *et al*., 2007). To determine if any of the three *CYCD3* genes are expressed during lateral shoot development, we used *CYCD3::GUS* reporter lines (Dewitte *et al*., 2007). No GUS activity was detected for *CYCD3;1* in any of the developmental stages of lateral shoots (data not shown), while both *CYCD3;2* and *CYCD3;3* promoters are active in SI and SII (Fig. 3C). *CYCD3;2* continues to be active throughout SIII, while *CYCD3;3* is switched off (Fig. 3C).

This shows that during the early phase lateral shoot cells are actively dividing and cell division contributes to the branch growth during SII and SIII. To understand if CK are involved in mediating the differences in cell proliferation between the adaxial and abaxial side of SII-SIII, mutants of the *ISOPENTENYLTRANSFERASES* (*IPT*) genes involved in CK biosynthesis were analysed (Takei, Sakakibara and Sugiyama, 2001). The *ipt3ipt5ipt7* triple mutant, containing diminished levels of CK (Miyawaki *et al*., 2006) shows reduced lateral shoot rootward growth during SII and SIII (Fig. 3D). To understand if the reduce rootward growth shown by *ipt3ipt5ipt7* is caused by alterations in cell proliferation along the ad-abaxial axis, number of cells in the triple mutant were analysed. *ipt3ipt5ipt7* shows reduced differences, with no statistical significance, in the number of cells between adaxial and abaxial side of SII-SIII lateral shoots compared to WT (Fig. 3E). This demonstrates that the phenotype shown by *ipt3ipt5ipt7* during lateral shoot early development is likely caused by attenuated differences in cell proliferation across the ad-abaxial axis.

### *TCP1* shows adaxial expression during lateral shoot development

To identify molecular regulators of the ad-abaxial polarity in lateral shoots during early development, we performed a comparative transcriptomic analysis of adaxial and abaxial sides of SII lateral shoots. We found that *TCP1*, a member of the *TEOSINTE BRANCHED1, CYCLOIDEA* and *PCF* (*TCP*) family, is predominantly expressed on the adaxial side of lateral shoots in both early and late stages of development (Fig. 4A), suggesting a possible role of TCP1 in the specification of adaxial identity. To confirm the role of TCP1 in regulating rootward growth in young lateral branches, we overexpressed *TCP1* coding sequence under the *CaMV 35S* constitutive promoter in WT plants (*p35S::TCP1*). *TCP1* overexpression shows pleiotropic effects in the shoot of T_1_ transgenic lines. Plants are overall smaller and have leaves with variable shapes (Fig. 4B). Interestingly, during lateral shoot development of *p35S::TCP1* the rootward growth of early stages is completely suppressed, and lateral shoots at SIV display very vertical GSAs similar to the primary shoot (Fig. 4C). Taken together these data suggest a possible role of TCP1 in the establishment of lateral shoot adaxial identity. Impairments in the ab-adaxial polarity could eventually lead to alteration of the final GSA displayed by lateral shoots in SIV.

**Figure 4:**
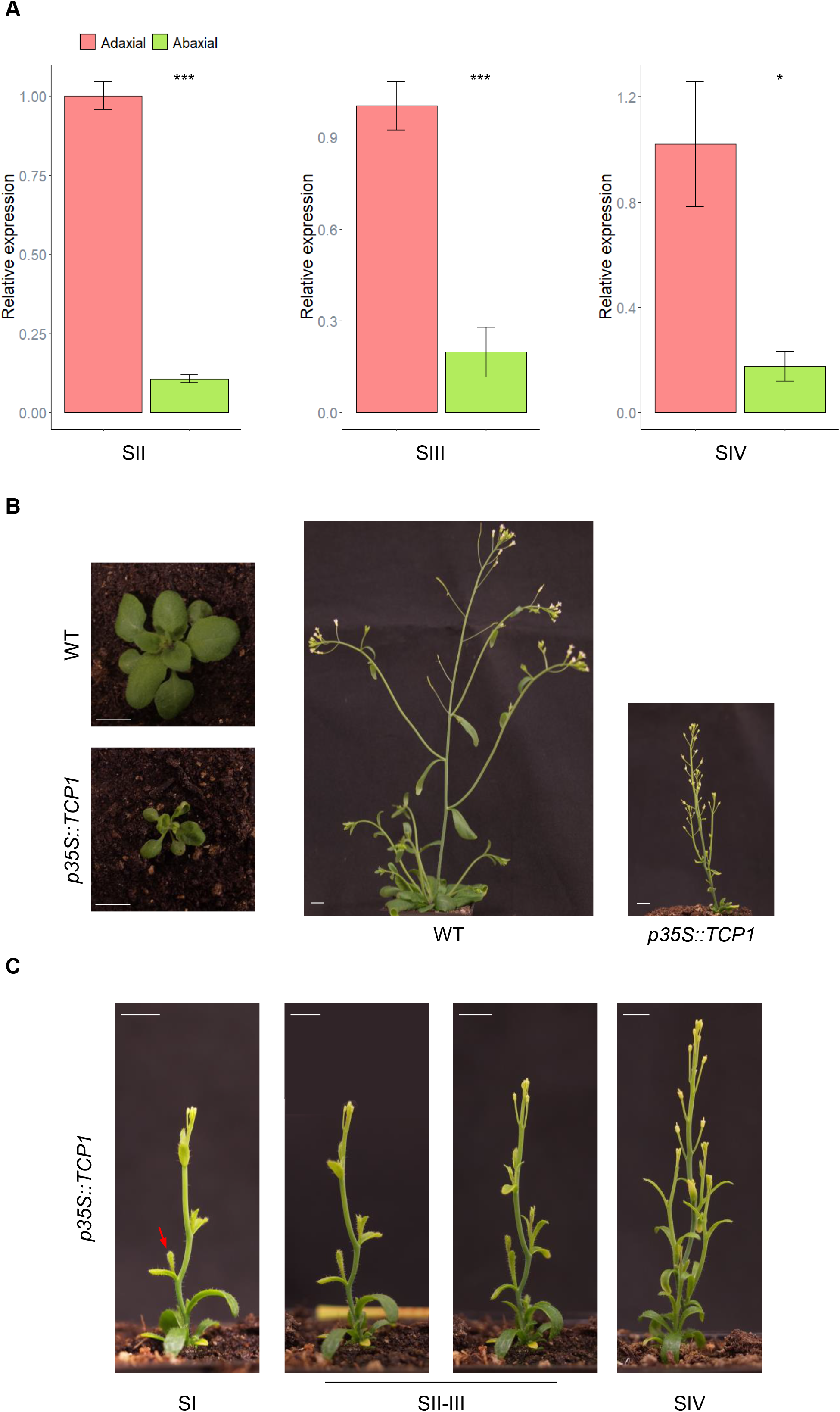
*TCP1* is expressed adaxially during lateral shoot development. A) Relative expression of *TCP1* on the adaxial and abaxial side of SII, SIII and SIV lateral shoots. Statistical analyses were performed using t-test. (*) = p < 0.05, (**) = p < 0.005. B) Representative images of *p35S::TCPI* (T_1_) and WT plants. Scale bars = 1 cm. C) Representative images of *p35S::TCPI* during lateral shoot development. Red arrow indicates SI lateral shoot. Scale bars = 0.5 cm.

## Discussion

Branching is a fundamental trait of shoot architecture and contributes to plant fitness and productivity. In both herbaceous and woody plants, the manipulation of shoot branching offers new opportunities for improvement of agronomically important species (Hill and Hollender, 2019). Multiple studies have been focused on understanding the regulation behind axillary meristems formation and activation leading to lateral shoot development, and the subsequent control of lateral shoot position with respect to gravity (Roychoudhry *et al*., 2013; Wang and Jiao, 2018). Here, we demonstrated the existence of a new developmental phase between the axillary meristem outgrowth and the setting of the GSA in mature branches. To accurately describe the events following axillary bud outgrowth, we divided lateral shoot development in four stages, with stages from I to III marking the early developmental phase and SIV the GSA setting. During these early stages lateral shoots do not maintain GSAs, but can perceive gravity and, to a certain extent, respond to gravistimulation. Our reorientations assays demonstrate that a simple tilting of the plant, which elicits a fast gravitropic response in SIV branches, is not enough to induce a tropic response in SII-III lateral shoots. Conversely, lateral shoots respond to gravistimulation if the reorientation determines a change in the branch polarity with respect to its plane of growth (Fig. S1). Our evidence suggests that ad-abaxial differences in cell proliferation are at the base of the rootward growth in SII, although at this stage, we cannot completely exclude the participation of cell elongation in this process. The heterogeneity of cell length in SII-SIII epidermis makes difficult to obtain any conclusive data regarding differences in cell elongation between the adaxial and abaxial side of young lateral shoots. The expression of D3 cyclins in lateral shoot stems from SI to SIII suggests that, at these stages, growth is sustained by cell proliferation. Although we observed ad-abaxial differences in cell number, we did not identify any asymmetry in *CYCD3* expression using GUS reporter lines. Analysis of other cell cycle regulators upstream and downstream of D-type cyclins, together with analysis of the protein levels of these regulators, that can be significantly different from the activity of their promoters, will elucidate the role of cell proliferation in marking ad-abaxial differences in young lateral shoots. Cytokinins have been shown to regulate cell cycle during leaf development through the induction of CYCD3 (Dewitte *et al*., 2007). In addition, CKs are involved in the positive regulation of both AM initiation and outgrowth (Waldie and Leyser, 2018). We found that mutants with reduced levels of CK display a reduction in SII rootward growth and cell proliferation differences between the adaxial and abaxial side of young lateral shoots. This suggests a possible involvement of CK in regulating cell proliferation during lateral shoot development. No differences in lateral shoot outgrowth kinetics were observed for the CK receptors *ARABIDOPSIS HISTIDINE KINASES* (*AHK*) loss-of-function mutants *ahk2ahk3, ahk2ahk4* and *ahk3ahk4* (data not shown), nor for the constitutively active gain-of-function variants of the AHK2 and AHK3 *repressor of cytokinin deficiency2* (*rock2*) and *rock3* (data not shown). The lack of phenotypes showed by AHK double loss-of-function and gain-of-function mutants could be due to redundancy of these genes in controlling shoot development. Further analysis of the expression and distribution of AHKs and other components of CK signalling are necessary to better understand the role of CK in lateral shoot development.

Based on these results we propose a model in which ad-abaxial polarity in lateral shoots is set early before or during stage I, independently from the gravitropic stimulus. The differences between adaxial and abaxial identity determine the branch to growth rootward during stage II (Fig. S2). At this stage the lateral shoot is already gravicompetent, but the auxin-dependent gravitropic machinery, that will lead to adjustments of the branch position with respect to gravity, is not yet able to overcome the developmental rootward growth of the young lateral shoot.

Our data indicates that rootward growth of lateral shoots during SII is determined principally by ad-abaxial differences in cell proliferation, while the GSA setting and maintenance relies on auxin-dependent regulation of cell elongation (Fig. S2). In this way, the exit from the cell cycle and the differentiation of lateral shoot cells during SIII leads to a switch from a gravity-independent growth, based on cell proliferation, to a gravity-dependent growth based on cell elongation. Previous work demonstrated the existence of an antigravitropic offset (AGO) acting to counteract underlying gravitropic response in the branch and producing gravity-dependent non-vertical growth (Roychoudhry *et al*., 2013). Our study indicates that the AGO is a distinct mechanism from the developmentally-driven rootward bending of SII lateral shoots. Whereas AGO activity is gravity- and auxin transport dependent, the analysis of early lateral shoot development suggests that the adaxial identity is set independently from gravity and is based on cell proliferation possibly driven by CK action.

Transcriptomic analysis revealed that *TCP1* is differentially expressed at the adaxial side of lateral shoots, making this gene a possible regulator of adaxial identity. Previous work in Arabidopsis, using a chimeric form of TCP1 fused with the SRDX transcriptional repressor motif, has linked TCP1 function to the control of cell elongation in leaf and stems (Koyama, Sato and Ohme-Takagi, 2010). In other works, TCP1 has been linked to brassinosteroid (BR) biosynthesis through the positive regulation of the biosynthetic BR gene *DWARF4* (*DWF4*) (Guo *et al*., 2010; Gao, Zhang and Li, 2015). In these studies, a *TCP1* overexpressing line, named *tcp1-1D*, shows elongated leaf petioles and stems (Guo *et al*., 2010). Finally, TCP1 has been also shown to be involved in the regulation of flower monosymmetry in *Iberis amara* (Busch and Zachgo, 2007; Busch, Horn and Zachgo, 2014). Here, *IaTCP1* expression on the adaxial side of the petals has been suggested to regulate cell proliferation (Busch, Horn and Zachgo, 2014). In addition, *IaTCP1* overexpression in Arabidopsis shows similar phenotypes to our *AtTCP1* overexpression T_1_ lines (Busch and Zachgo, 2007). These data suggest that TCP1 may be involved in multiple roles during shoot development, regulating cell proliferation and/or cell elongation. Quantification of *TCP1* expression levels in our T_1_ *p35S::TCP1* lines and the analysis of the cell morphology and number in lateral shoots will further elucidate the role of TCP1 during branch development.

