## Supplemental figures for "Arabidopsis lateral shoots display two distinct phases of growth angle control"

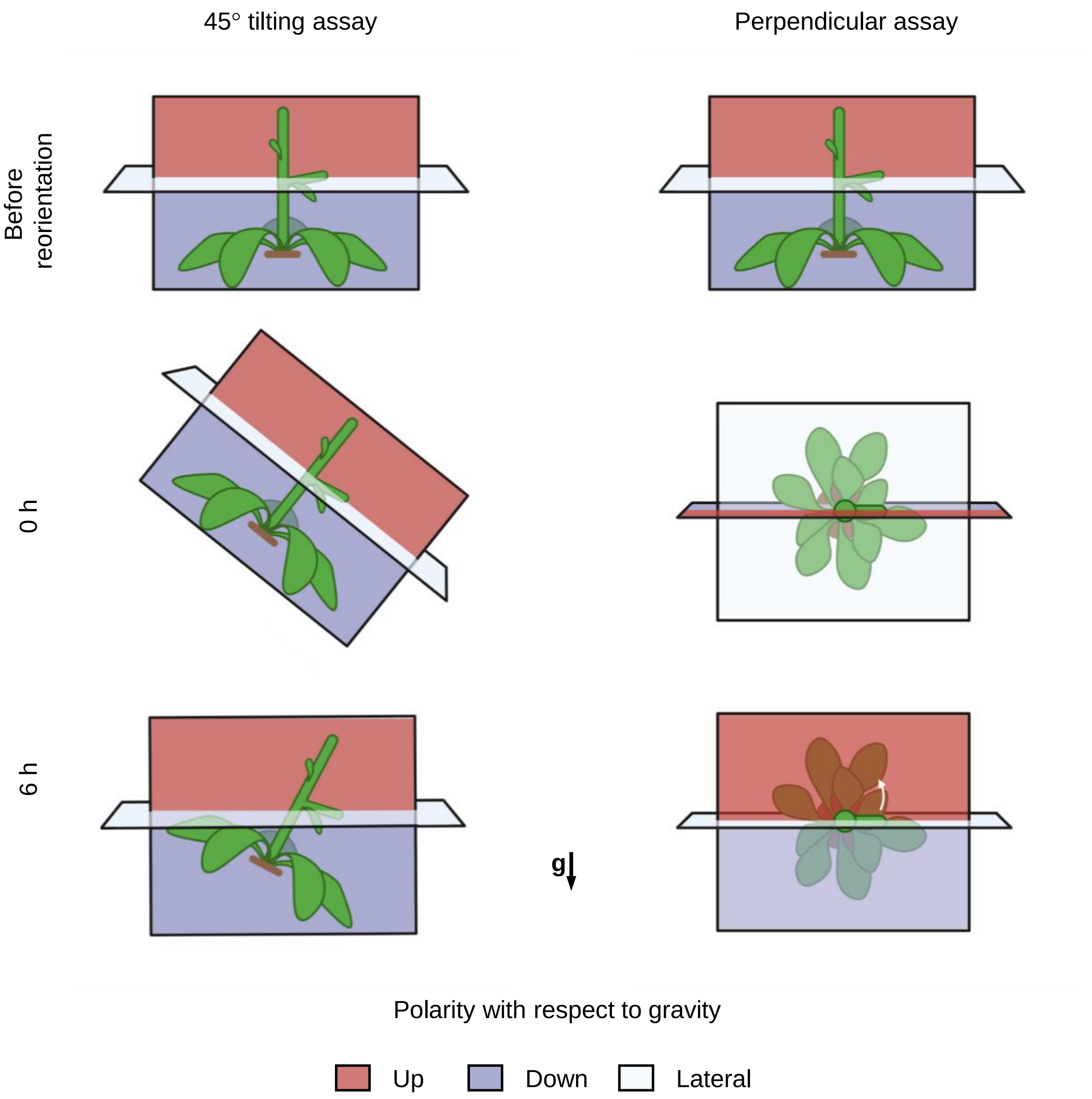


**Figure S1: Reorientation assays**
Schematic representation of 45° tilting and perpendicular assays. Black arrow indicates the direction of gravity, g = gravity.

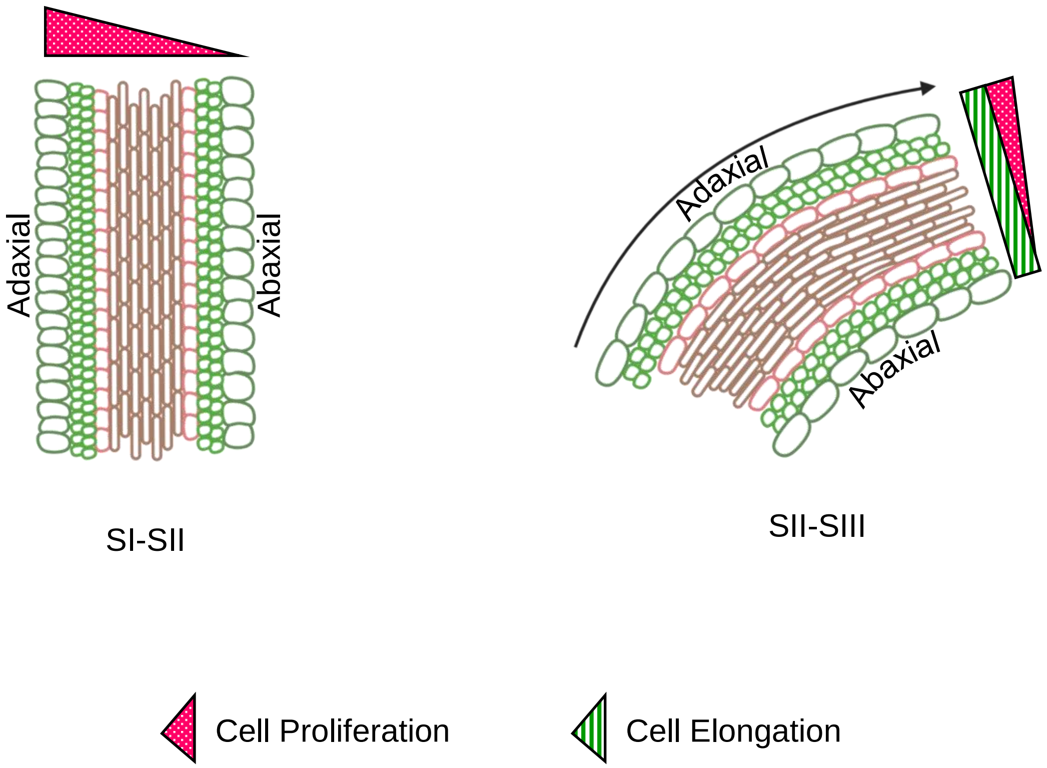


**Figure S2: Cell proliferation sustains early lateral shoot development**
Schematization of cell proliferation and cell elongation along the ad-abaxial axis during lateral shoot early development. Triangles indicate the magnitude of the processes. Black arrows indicate the direction of growth. During SI-SII greater cell proliferation at the adaxial side determines lateral shoot early rootward growth, while cell elongation happens symmetrically across the branch.
